# Pleasurable music activates cerebral μ-opioid receptors: A combined PET-fMRI study

**DOI:** 10.1101/2024.04.10.588805

**Authors:** Vesa Putkinen, Kerttu Seppälä, Harri Harju, Jussi Hirvonen, Henry K. Karlsson, Lauri Nummenmaa

## Abstract

The μ-opioid receptor (MOR) system mediates incentive motivation and the hedonic component of primary rewards such as food and sex. However, there is no direct *in vivo* evidence for the involvement of the MOR system in pleasure derived from aesthetic rewards such as music. We measured MOR activation with positron emission tomography (PET) and the agonist radioligand [^11^C] carfentanil with high affinity for MORs during the listening of pleasurable music and neutral baseline condition. Haemodynamic responses to pleasurable music were measured using functional magnetic resonance imaging (fMRI). The PET results revealed that pleasurable music increased [^11^C]carfentanil binding in several cortical and subcortical regions, including ventral striatum and orbitofrontal cortex, known to contain “hedonic hotspots”. Individual variation in baseline MOR tone influenced pleasure-dependent haemodynamic responses during music listening in regions associated with interoceptive, sensorimotor, and reward processing. Our results provide the first-ever neuroimaging evidence that listening to pleasurable music modulates MOR system activation and indicate that the μ-opioid system governs complex aesthetic rewards in addition to biologically salient primary rewards.

## Introduction

Pleasure motivates humans to seek out rewards crucial for survival and reproduction. Yet, human hedonic responses extend beyond such primary rewards and encompass abstract, aesthetic rewards that seemingly lack adaptive function. A prime example is music, which has persisted throughout human evolution and remains one of the most common sources of pleasure today despite offering no obvious survival advantage (however, see Mehr et al., 2021 and Savage et al., 2021). Nevertheless, neuroimaging studies indicate that music-induced pleasure engages the same hedonic circuitry as biologically salient primary rewards, including regions such as the ventral striatum, orbitofrontal cortex, and insula (Koelsch, 2014; Mas-Herrero et al., 2021; Nummenmaa et al., 2021). However, the majority of these studies have been conducted using functional magnetic resonance imaging (fMRI), which is unable to resolve the underlying neurochemistry (however, see Salimpoor et al., 2011). Given the importance of specific neuromodulator systems in governing human emotion (Fox & Lobo, 2019; Nummenmaa & Tuominen, 2017), unraveling the molecular neural basis of musical hedonia is crucial for understanding the universal human affinity towards music.

The μ-opioid receptor (MOR) system plays a central role in incentive motivation and the hedonic components of primary rewards. Studies in rodents indicate that subregions in the nucleus accumbens, ventral pallidum, insula, and orbitofrontal cortex contain cell assemblies whose activity is causally related to hedonic reactions (Berridge & Kringelbach, 2015). Injection of opioids into these regions heightens liking behaviors, indicating that MORs mediate pleasure associated with reward consumption (e.g., Peciña & Berridge, 2005). In humans, opioid agonists and antagonists modulate the hedonic impact of rewards like sweet taste (Eikemo et al., 2016) and sexual stimuli (Buchel et al., 2018). Multivariate pattern analysis of fMRI data has revealed that activity associated with subjective pleasure overlaps with regions showing mu-opioid receptor gene expression and is down-regulated by an opioid antagonist (Kragel et al., 2023). Positron emission tomography (PET) studies using the MOR-specific radioligand [^11^C]carfentanil have revealed opioid release following biologically salient rewards such as eating and sex (Jern et al., 2023; Manninen et al., 2017), and baseline MOR availability modulates heamodynamic and behavioral responses to emotional stimuli (Manninen et al., 2017; Sun et al., 2022). Although the contribution of the MOR system musical hedonia has also been postulated (Nummenmaa et al., 2021; Tarr et al., 2014), this hypothesis remains to be tested with in vivo neuroimaging.

Several lines of evidence suggest, however, that the MOR system might be engaged during musical hedonia. First, given that the MOR system mediates analgesia, the pain-relieving and prosocial effects of pleasurable music lend indirect support for opioidergic contribution to music-induced pleasure (Nummenmaa et al., 2021; Tarr et al., 2014): Listening to pleasurable music alleviates chronic and post-operative pain and reduces the need for opioid pain medication in patients (Kühlmann et al., 2018; Sihvonen et al., 2022) while engaging in musical activities such as singing and dancing heightens pain threshold in healthy populations (Dunbar et al., 2012). Secondly, musical activities foster social bonding and promote cooperation (Dunbar et al., 2012; Kirschner & Tomasello, 2010; Pearce et al., 2015), and the involvement of opioids in affiliative behavior, established through pharmacological interventions in non-human primates (Fabre-Nys et al., 1982; Graves et al., 2002) and PET studies in humans (Manninen et al., 2017; Nummenmaa et al., 2016), suggests that the MOR systems may mediate these prosocial effects of music. Finally, pharmacological down- and up-regulation of endogenous opioid function have yielded partial support for opioidergic contribution to music-induced pleasure. The earliest study (Mallik et al., 2017) observed diminished music-induced pleasure after opioid receptor blockade with naltrexone. Two subsequent pharmacological studies, however, failed to replicate the effect on subjective pleasure but found that opioid agonist and/or antagonist-induced modulation of neurophysiological markers of music-induced arousal (Laeng et al., 2021; Mas-Herrero et al., 2023).

### The current study

To directly test whether musical pleasures engage the MOR system, we measured MOR activation *in vivo* with PET using the high-affinity agonist radioligand [^11^C]carfentanil while participants listened to self-selected pleasurable music versus a neutral control condition. We also measured haemodynamic responses to the same musical excerpts in a separate fMRI experiment to unravel the interplay between the MOR system and the acute pleasure-dependent haemodynamic activity. We also quantified autonomic nervous system activity during music listening by measuring heart rate and pupil size. We found that i) pleasurable music increased MOR availability relative to a neutral baseline condition in regions associated with emotion and reward (e.g., ventral striatum, amygdala, orbitofrontal cortex) and ii) elicited higher pleasure-dependent haemodynamic responses in the auditory, somatosensory, and emotion networks which were stronger in participants with higher baseline MOR availability. The heart rate and pupil size measurements indicated that pleasurable music induced heightened autonomic arousal. These data indicate that the MOR system significantly contributes to musical pleasure and that individual variation in MOR availability may constitute a molecular mechanism explaining individual differences in the propensity for enjoying music.

## Methods

### Participants

Fifteen women (mean age 26.0 years, range 19-42) participated in both the PET and fMRI experiments. Fifteen additional women (mean age 23.33 years, range 19-27) participated in the fMRI. All the fMRI-only participants as well as eighteen additional women (mean age 25.7 years, range 20-37), participated in the eye-tracking experiment. The participants were recruited through university email lists. To ensure that the participants would enjoy listening to music, potential participants filled in the Emotional Evocation, Mood Regulation, and Social Reward facets of the Barcelona Music Reward Questionnaire (BMRQ) (Mas-Herrero et al., 2013), and those who received at least 80% of the possible maximum score were invited to participate. Only women were studied to maximize statistical power due to sex-dependent variability in the spatial distribution of MORs (Kantonen et al., 2020). Additionally, women report stronger emotional responses (Lench et al., 2011), thus female-only sample was assumed to maximize the effects in the complex PET studies. The study physician screened the participants for eligibility, and a psychologist screened them for psychiatric disorders with the M.I.N.I 6.0 interview (Sheehan et al., 1998). The exclusion criteria included a history of neurological or psychiatric disorders, alcohol and substance abuse, current use of medication affecting the central nervous system, and the standard MRI exclusion criteria. Structural brain abnormalities that are clinically relevant or could bias the analyses were excluded by a consultant neuroradiologist. All participants gave informed, written consent and were compensated for their participation. The ethics board of the Hospital District of Southwest Finland had approved the protocol, and the study was conducted in accordance with the Declaration of Helsinki.

### Stimuli

We used self-selected music as stimuli to maximize their emotional impact (Salimpoor et al., 2011). Each subject compiled an approximately 90-min playlist of music that elicited strong pleasure in them. Most stimuli were contemporary pop, R&B, and rap/hip-hop (**See Figure S1**).

### PET experimental protocol

Participants took part in two PET scans: a music (“challenge”) scan and a neutral baseline scan. Before the music scan, the subject listened to their playlist through headphones alone for 15 minutes. During the PET acquisition, the participants lay in the PET scanner wearing hospital clothes and continued to listen to their playlist through headphones. The baseline scan was otherwise identical but without the musical stimulation. During both scans, the participants were asked to verbally report their level of felt pleasure on a scale of 1-10 (no pleasure to high pleasure) every 10 minutes. The participants also reported the occurrence of music-induced chills with a button box. The music and baseline scans were completed on separate days at the same time of the day, and the order of the scans was counterbalanced across participants.

### PET data acquisition

MOR availability was measured with radioligand [^11^C]carfentanil synthesized as described previously (Hirvonen et al., 2009). The radiochemical purity of the produced [^11^C]carfentanil batches was 98.1 ± 0.4 % (mean ± SD). The injected [^11^C]carfentanil radioactivity was 252 ± 10 MBq, and molar radioactivity at the time of injection was 354 ± 240 MBq/nmol, corresponding to an injected mass of 0.52 ± 0.43 μg. PET imaging was carried out with Discovery 690 PET/CT scanner (GE Healthcare, US). The tracer was administered as a single bolus via a catheter placed in the participant’s antecubital vein, and radioactivity was monitored for 51 minutes. The participant’s head was strapped to the scan table to prevent excessive head movement. T1-weighted MR scans were obtained for attenuation correction and anatomical reference.

### PET image processing and data analysis

PET data were preprocessed with the Magia (Karjalainen et al., 2020) toolbox (https://github.com/tkkarjal/magia) running on MATLAB (The MathWorks, Inc., Natick, MA, USA). PET images were first motion-corrected and coregistered to T1-weighted (T1w) MR images, after which T1w image was processed with Freesurfer for anatomical parcellation. [^11^C]carfentanil uptake was quantified as the binding potential (*BP*_ND_) relative to non-displaceable binding, estimated with the simplified reference tissue model (SRTM) at voxel-level by using the occipital cortex as the reference region. *BP*_ND_ images were spatially normalized to MNI152-space and smoothed using a Gaussian kernel (FWHM = 6 mm).

The voxel-vise differences between the music and baseline conditions in MOR availability were assessed in SPM12 (http://www.fil.ion.ucl.ac.uk/spm/) using a repeated-measures t-test. The statistical threshold was set at p > 0.05, FWE-corrected at the cluster level.

### fMRI experimental protocol

The fMRI experiment was run on a separate day after the PET scans. The participants were presented with ten 45-sec segments extracted from their self-selected pleasurable musical pieces (see above). They were also presented with two 45-sec random tone sequences as control stimuli. Stimuli were presented binaurally via MRI-compatible headphones (Sensimetrics S14) at a comfortable level adjusted individually for each participant. Participants were asked to remain still during the scan and focus on the feelings evoked by the music. The subject used a button box to move a cursor on the screen to indicate the current music-induced pleasure from 0 (no pleasure) to 10 (high pleasure) during the music and the control stimulation blocks. The starting position of the cursor was randomized. The stimulation blocks were presented in a random order and were interspersed with 30 second rest blocks with no stimulation.

### MRI data acquisition and preprocessing

The MRI data were acquired using a 3T MRI system with SuperG gradient technology (SIGNA, Premier, GE Healthcare, Waukesha, WI, USA) with the 48-channel head coil. High-resolution structural images were obtained with a T1-weighted (T1w) MPRAGE sequence (1 mm^3^ resolution, TR 7.3 ms, TE 3.0 ms, flip angle 8°, 256 mm FOV, 256 × 256 reconstruction matrix). 556 functional volumes (24 min) were acquired with a T2∗-weighted echo-planar imaging sequence sensitive to the blood-oxygen-level-dependent (BOLD) signal contrast (TR 2600 ms, TE 30 ms, 75º flip angle, 240 mm FOV, 80 × 80 reconstruction matrix, 3.0 mm slice thickness, 45 interleaved axial slices acquired in descending order without gaps).

Functional imaging data were preprocessed with FMRIPREP. During preprocessing, each T1w volume was corrected for intensity non-uniformity using N4BiasFieldCorrection (v2.1.0) and skull-stripped using antsBrainExtraction.sh (v2.1.0) using the OASIS template. Brain surfaces were reconstructed using recon-all from FreeSurfer (v6.0.1), and the brain mask estimated previously was refined with a custom variation of the method to reconcile ANTs-derived and FreeSurfer-derived segmentations of the cortical grey matter of Mindboggle. Spatial normalization to the ICBM 152 Nonlinear Asymmetrical template version 2009c was performed through nonlinear registration with the antsRegistration (ANTs v2.1.0), using brain-extracted versions of both T1w volume and template. Brain tissue segmentation of cerebrospinal fluid, white matter and grey matter was performed on the brain-extracted T1w image using FAST (FSL v5.0.9).

### Regional Effects in the General Linear Model

The fMRI data were analyzed in SPM12 (Wellcome Trust Center for Imaging, London, UK, (http://www.fil.ion.ucl.ac.uk/spm). To reveal regions whose activity correlated with music-induced pleasure, a general linear model (GLM) was fit to the data where the design matrix, including a boxcar regressor for the music block vs. silent periods with subjective pleasure rating as a parametric modulator of interest (cf. Martínez-Molina et al., 2016). A stick function nuisance regressor for the button presses was also included. For each subject, contrast images were generated for the main effect of pleasure. The contrast images were then subjected to a second-level analysis for population-level inference. Clusters surviving family-wise error rate (FWE) correction (p < 0.05) are reported.

### PET-MRI fusion analysis

To examine the connection between baseline MOR tone and hemodynamic responses in music-induced pleasure, we calculated mean subject-wise baseline MOR availabilities across the 17 ROIs (amygdala, caudate, cerebellum, dorsal anterior cingulate cortex, inferior temporal cortex, insula, middle temporal cortex, nucleus accumbens, orbitofrontal cortex, pars opercularis, posterior cingulate cortex, putamen, rostral anterior cingulate cortex, superior frontal gyrus, superior temporal sulcus, temporal pole, and thalamus) defined by the FreeSurfer parcellations. We then correlated the ROI-wise MOR availabilities with the regional pleasure-dependent BOLD responses. Finally, we generated a cumulative map of MOR-dependent BOLD responses to illustrate the brain regions where haemodynamic activity was consistently associated with MOR availability.

### Heart rate and pupil size measurements

We assessed autonomic nervous system activation by comparing mean heart rate during the music and baseline PET scans. Heart rate was measured using a Polar M430 GPS running watch, and a Polar H10 heart rate sensor.

In the eye-tracking experiment, we measured pupil size with Eye Link II system with 250 Hz sampling rate and spatial accuracy better than 0.5 degrees. The recording was conducted in a dimly lit room. The participants were seated with their chin on a chin rest. The eye tracker was calibrated and validated using standard 9-point calibration. The participants listened to 10 60-sec excerpts of their self-chosen pleasurable music and six 45-sec control stimuli used in the fMRI experiment (see above). A control stimulus was presented after 2-3 music excerpts. Each trial began with a drift correction and detrending. Participants were instructed to keep their eyes fixated at a cross shown at the center of the screen while listening. Gaze position and pupil size were measured throughout the trial, after which the subject reported using a keyboard their experience of liking, calmness, and feeling energized on a scale from 1 to 4 (1 = very weak, 4 = very strong) (**Figure S2**). The eye tracker was recalibrated at the middle of the experiment. Subject-wise pupil size time series were cleaned from blinks using in-house code based on PhysioData Toolbox (Kret & Sjak-Shie, 2019), baseline corrected (0-20 ms), and mean pupil sizes between 2 and 10 seconds was compared across the music and control trials.

## Results

### Subjective pleasure evoked by music

We analyzed pleasure ratings obtained during the PET scans with a linear mixed-effect model with Time (0, 10, 20, 30, 40 min) and Condition (music vs. baseline) as fixed factors and subject as a random factor (pleasure ∼ time * condition + (1|subject)). Participants experienced significantly more pleasure during the music scan than during the baseline scan (Main effects of Condition: F(1, 32) = 45.135, *p* < .001, **Figure 1a**). The participants reported an average of 6 (SD = 4.80) chills during the music scan. We extracted the maximum pleasure rating for the music and control blocks from the continuous ratings obtained during the fMRI scan and then calculated separate averages for the music and control conditions for each subject. A repeated measures t-test indicated that the ratings were higher for the music than for the control condition (t(32) = 28.504, p < .001) (**Figure 1b**).

**Figure 1.**
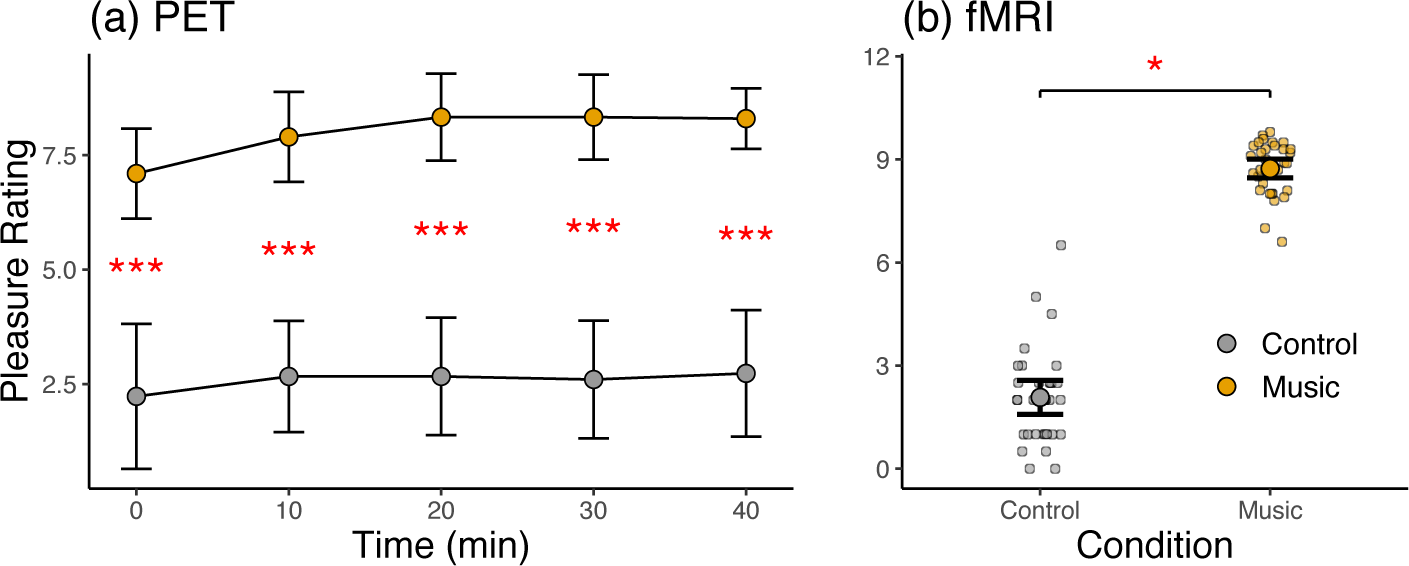
Pleasure ratings during the PET scans for the music and control conditions (a) and the music and control blocks in fMRI (b). The error bars show 95% confidence intervals. *** = *p* < .001, * = *p* < .05

### Eye tracking and heart rate

In the eye-tracking experiment, the participants’ pupil size was significantly larger for the music excerpts than for the control stimuli between 2-10 sec from stimulus onset (*t*(32) = 3.268, *p* < .01.) (**Figure 2a and 2b**). During the PET scans, the participants’ mean heart rate was higher for the Music than for the Baseline scans (*t*(7) = 4.104, *p* < .01.) (**Figure 2c**).

**Figure 2.**
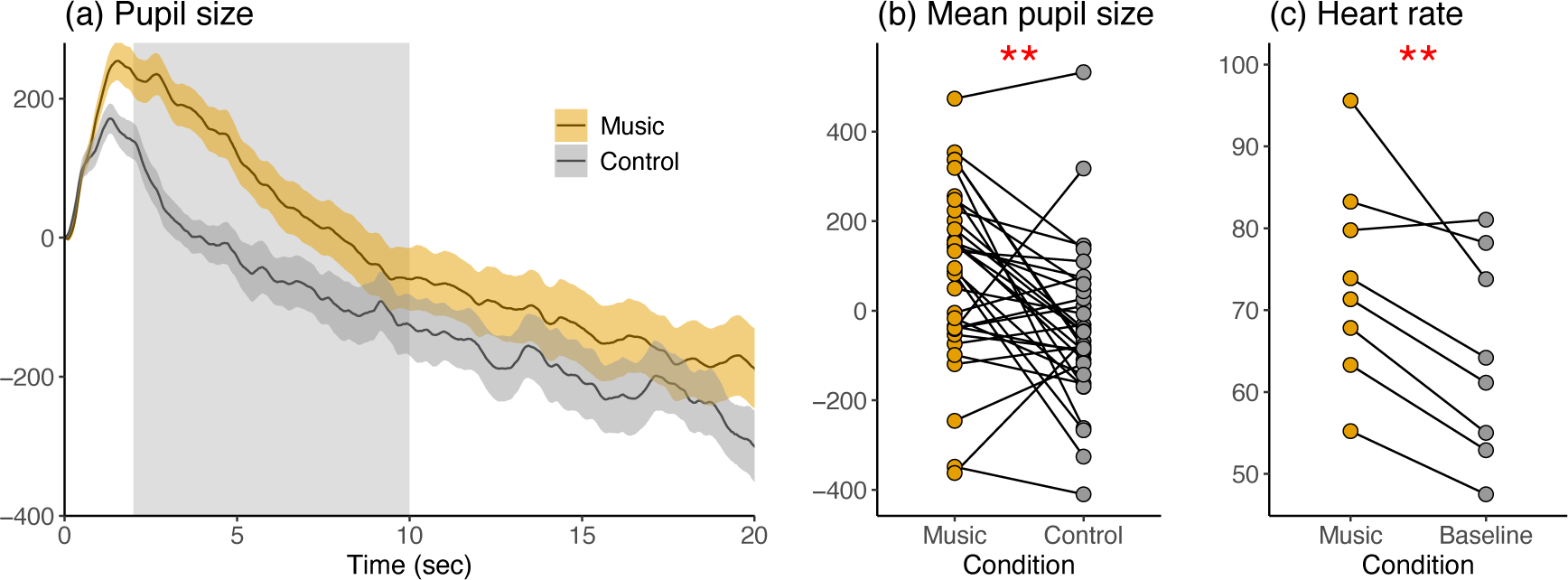
(a) Average pupil size as a function of time for the pleasurable music experts and neutral control stimuli during the eye-tracking experiment. The shaded area around the curves indicates 95% confidence interval. The shaded rectangle indicates the time window (2-10 sec) used to quantify mean pupil size (b) The mean pupil size per subject for the music and control stimuli (c) The mean heart rate during the Music and Baseline PET scans. ** = *p* < .01.

### PET

Whole-brain analysis of the PET data revealed higher *BP*_ND_ in the music condition relative to the control condition in several brain regions, including ventral striatum, amygdala, parahippocampal gyrus, thalamus, brainstem, orbitofrontal cortex, and temporal pole (**Figure 3**). The opposite contrast did not reveal significant effects.

**Figure 3.**
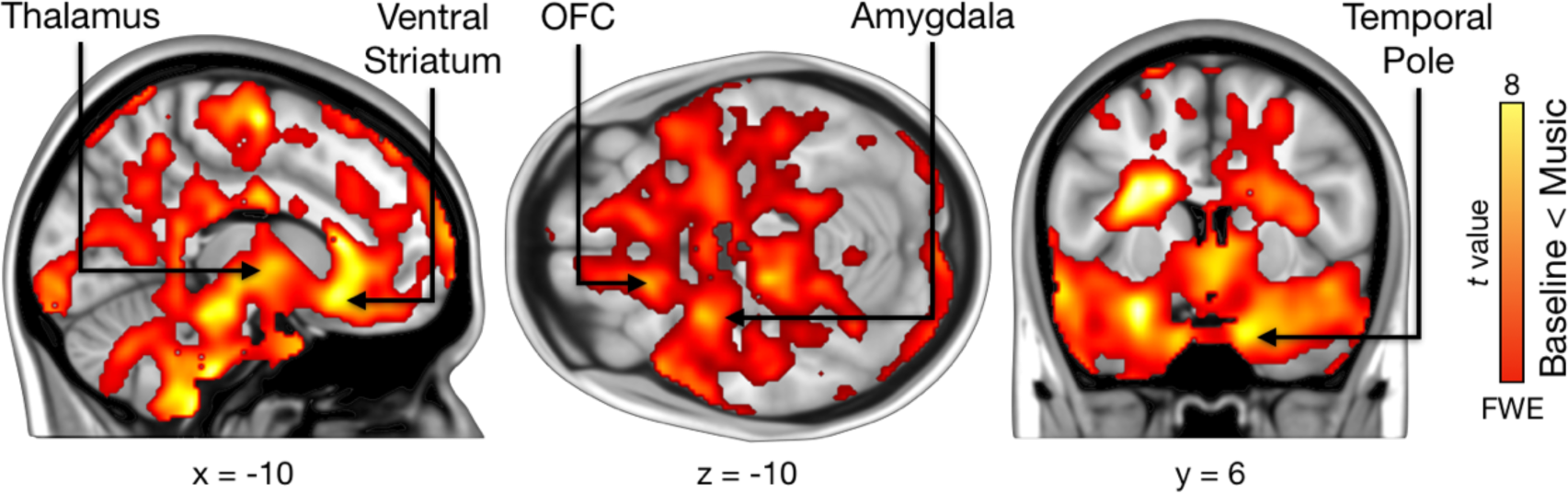
Regions showing increased *BP*_ND_ during music listening relative to neutral baseline. The data are thresholded at *p* < .05 and FWE corrected at the cluster level. Colourbar indicates the t statistic range.

### fMRI

Modeling the BOLD data with the parametric pleasure rating revealed activity in the insula, orbitofrontal cortex (OFC), and ACC, as well as in the middle frontal gyrus and frontal pole. Subcortically, activation was observed particularly in right caudate and bilateral putamen. Activation was also seen in sensorimotor regions in the pre and postcentral gyri, SMA, and supramarginal gyrus as well as in the visual cortex (**Figure 4**).

**Figure 4.**
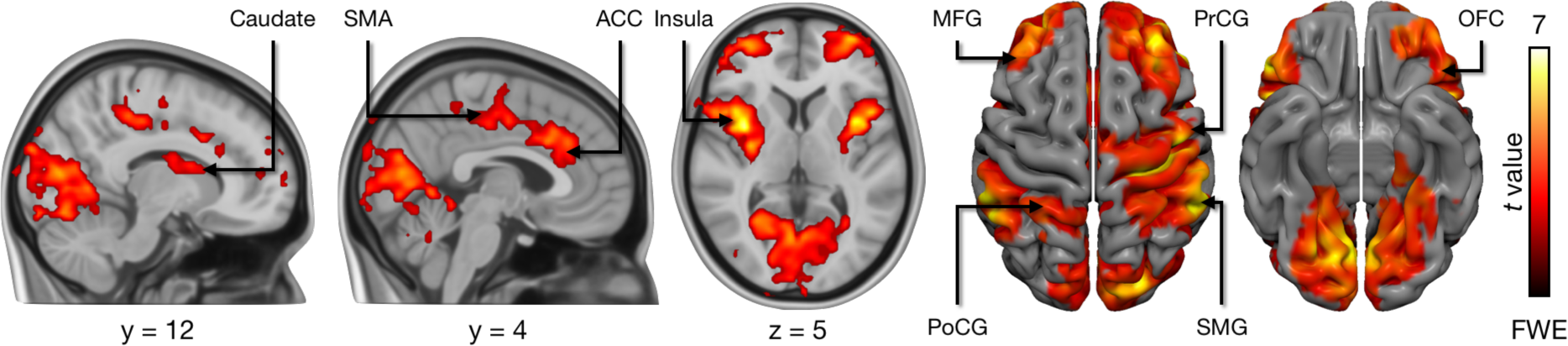
Pleasure-dependent BOLD responses. The data are thresholded at *p* < .05 and FWE corrected at cluster level. ACC = Anterior cingulate cortex, MFG = Middle frontal gyrus, PoCG = Postcentral gyrus, PrCG = precentral gyrus, SMA = Supplementary motor area, SMG = supramarginal gyrus, OFC = Orbitofrontal gyrus. Colourbar indicates the t statistic range.

**Figure 5.**
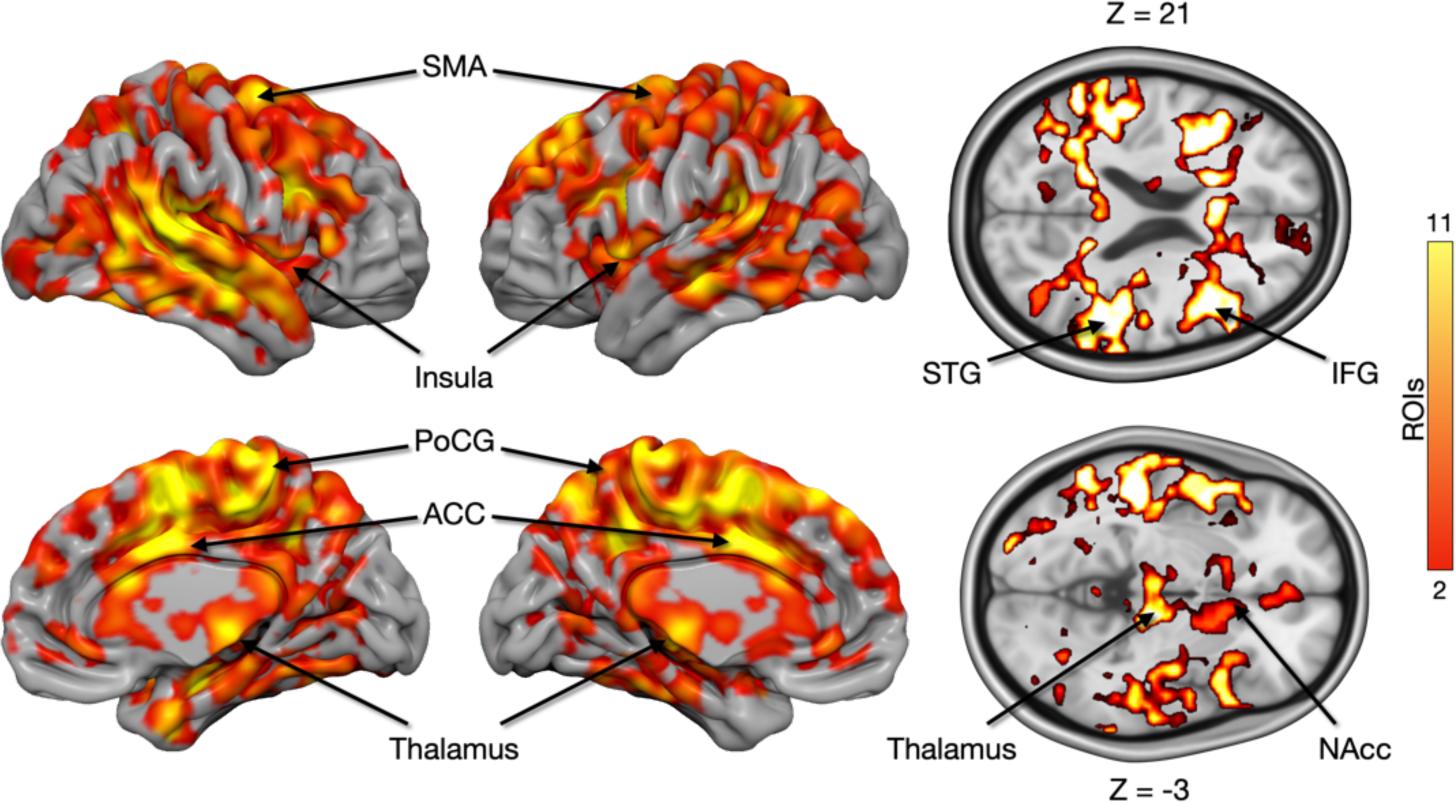
No significant negative associations between MOR tone and haemodynamic responses were found. Cumulative maps showing the number of ROIs (out of 17) where BP_ND_ was positively correlated with pleasure-dependent BOLD responses. ACC = Anterior cingulate cortex, PoCG = Postcentral gyrus, SMA = Supplementary motor area.

### PET-fMRI fusion analysis

Finally, we tested whether regional baseline MOR availability in 17 a priori selected ROIs was associated with the whole brain pleasure-dependent BOLD responses. This analysis revealed significant positive associations with the regional *BP*_ND_ and BOLD responses in the Insula, ACC, SMA, post central gyrus, and middle and superior temporal gyrus, ventral striatum, and thalamus

## Discussion

Our main finding was that pleasurable music activated the MOR system as evidenced by higher [^11^C]carfentanil binding in several cortical and subcortical regions, including the reward circuit. The fMRI results revealed that acutely music-induced pleasure was associated with activity in regions implicated in interoception, emotion, and reward. Finally, the PET-fMRI fusion analysis demonstrated that higher baseline MOR availability was associated with stronger pleasure-dependent BOLD responses particularly in the reward circuit as well as in motor regions. Altogether, our results represent the first *in vivo* neuroimaging evidence supporting the mediating role of the MOR system in the experience of music-induced pleasure and underline how interindividual variability in the MOR system is associated with individual differences in the tendency for reward responses to music.

### Music modulates MOR activation

The PET data revealed that pleasurable music modulated opioidergic activity in central nodes of the reward circuits such as the ventral striatum and OFC containing “hedonic hotspots” regulated by opioids (Berridge & Kringelbach, 2015). While prior [^11^C]carfentanil PET studies on reward processing have mainly focused on biologically salient primary rewards such as feeding (Tuulari et al., 2017), sex (Jern et al., 2023), and sociability (Manninen et al., 2017), our results indicate that aesthetic rewards, such as music, that are strongly influenced by cultural learning, also modulate MOR system activity. This suggests that MORs mediate pleasure across domains, driving humans to pursue primary and non-primary rewards and make choices in the face of conflicting options (Grabenhorst & Rolls, 2011). Due to the well-established role of endogenous opioids in affiliative behavior and social rewards, the MOR system has been hypothesized to contribute particularly to the prosocial effects of musical activities involving synchronized movements in groups, such as joint music-making and dancing (Dunbar et al., 2012; Savage et al., 2021). Our results, however, indicate that even solitary listening to pleasurable music without overt movement can modulate MOR system activity.

We observed higher *BP*_ND_ in the music condition compared to the control condition. A decrease in *BP*_ND_ is typically interpreted as evidence for heightened endogenous neurotransmitter release, in line with competition between the radiotracer and synaptic neurotransmitters (Colasanti et al., 2012; Mick et al., 2014), while an elevation in [^11^C]carfentanil binding may signify MOR “deactivation”, a reduction in synaptic endogenous opioids (Nummenmaa et al., 2016). However, *BP*_ND_ encompasses both receptor density and affinity, and augmented radioligand binding may indicate an increase in the available receptors or enhanced binding affinity. While prior studies indicate that pleasurable stimuli often decreases [^11^C]carfentanil *BP*_ND_, pleasurable touch, laughter, and social acceptance (Hsu et al., 2013; Manninen et al., 2017; Nummenmaa et al., 2016) have also been reported to heighten regional MOR availability. The current design does not allow us to disentangle whether the observed effects are due to increased or decreased opioid tone. Even though the exact cellular mechanism cannot be resolved due to the limitations of the PET technique, our findings support the overall engagement of the MOR system during the experience of music-induced pleasure.

While endogenous opioids have often been hypothesized to mediate musical hedonia (Chanda & Levitin, 2013; Nummenmaa et al., 2021; Tarr et al., 2014), an alternative proposal is that the dopamine system constitutes the main neurochemical pathway for music-induced pleasure (Mas-Herrero et al., 2023). This assertion is supported by a PET study demonstrating heightened dopamine release during music-induced chills (Salimpoor et al., 2011) and pharmacological data showing that dopamine receptor blockade diminishes music-induced pleasure and dopamine agonist administration had the opposite effect (Ferreri et al., 2019). Animal studies have, however, demonstrated that injecting opioid agonists into the striatum increases liking reactions, whereas dopamine antagonist injections in these sites or chemical (6-OHDA) lesions of the striatal dopaminergic neurons do not reduce liking responses (Berridge et al., 1989; Berridge & Kringelbach, 2015). Moreover, one human study has reported opioid antagonist-dependent weakening of pleasure induced by music (Mallik et al., 2017), yet this outcome has not been replicated in subsequent research (Laeng et al., 2021; Mas-Herrero et al., 2023). The dopamine and opioid systems interact at the molecular level (Tuominen et al., 2015), and it is thus likely that the interaction between the systems plays a role in shaping the experience of music-induced pleasure and arousal (Chanda & Levitin, 2013).

Opioidergic contribution to music-induced pleasure also has translational implications. Music-based interventions aid post-operative patient recovery by reducing pain, anxiety, and the need for analgesic medication (Hole et al., 2015), and the present data suggest that MORs may mediate these effects. Depressive symptomology and anxiety, which are prominent symptoms in many psychiatric disorders, are associated with altered MOR availability (Nummenmaa et al., 2020). Music therapy, effective in treating depression and other mental disorders (Gold et al., 2009), may thus partly rely on opioid influence. Music-based interventions show promise in non-invasive rehabilitation for Parkinson’s disease, dementia, stroke, acquired brain injuries, and neurological disorders (Sihvonen et al., 2017). MORs may contribute to the efficacy of these interventions through modulation of motivation and cognitive control (van Steenbergen et al., 2019).

### MOR tone modulates haemodynamic responses to pleasurable music

Our fMRI results revealed that activity in the orbitofrontal cortex and dorsal striatum correlated with the subjective experience of music-induced pleasure. Prior research has consistently shown that the OFC activity tracks subjective pleasure derived from various rewarding stimuli, such as food and sexual cues (Berridge & Kringelbach, 2015). In the dorsal striatum, the caudate and putamen are consistently activated by different rewards (Mas-Herrero et al., 2021), including liked music (Brattico et al., 2015; Martínez-Molina et al., 2016; Putkinen et al., 2021). Pleasure-dependent activation was also observed in the ACC and insula (Blood & Zatorre, 2001; Putkinen et al., 2021), which are associated with processing visceral signals and may contribute to the physiological arousal accompanying music-induced pleasure (Ferraro et al., 2022; Salimpoor et al., 2009). Accordingly, we found that pleasurable music elicited increased heart rate and stronger pupillary responses, indicating increased autonomic activity in line with previous studies showing that music-induced pleasure and emotional arousal coincide with heightened autonomic arousal (Koelsch & Jäncke, 2015). In particular, the insula is consistently activated by both music and food rewards (Mas-Herrero et al., 2021), possibly owing to its role in emotional and bodily responses that are central to music. We also found activation in the right pre and postcentral gyri, along with the supramarginal gyrus, further indicating the experience of music-induced pleasure relies on somatomotor system activation that may might mirror bodily sensations and simulations of movement triggered by liked music (Gordon et al., 2018; Putkinen et al., 2024).The pleasure-dependent BOLD responses co-localized with music-induced *BP*_ND_ changes in the insula, ACC, and frontal pole, suggesting that local blood flow increase in these regions during music-induced pleasure may partially reflect the metabolic needs of the MOR system.

Baseline MOR availability was associated with hemodynamic pleasure-dependent responses: Participants with a higher concentration of MORs exhibited stronger haemodynamic pleasure responses, particularly in the ACC, insula, and auditory cortex. MOR tone was also associated with activity in NAcc, which is a central node in the reward circuit of the brain and shows consistent activation in response to pleasurable music (Mas-Herrero et al., 2021). Thus, our results indicate that individual variation in MOR tone influences pleasure responses in regions associated with bodily, auditory, and reward processing during pleasurable music listening, which may explain individual differences in subjective music-induced emotional experience. It is noteworthy, however, that prior studies have shown that baseline MOR availability is associated with functional BOLD responses to both positive, reward-related stimuli, such as food pictures and laughter sounds, as well as negative stimuli, like violent videos (Karjalainen et al., 2017; Nummenmaa et al., 2018; Sun et al., 2022). MOR availability has also been associated with haemodynamics responses reflecting emotional arousal irrespective of valence. Together, these findings suggest the endogenous opioid system has a highly general role in modulating the processing of a wide range of emotionally evocative events irrespective of valence and biological saliency.

### Limitations

Due to the complexity of PET neuroreceptor imaging, our sample size was relatively small, which may have compromised the ability to detect small effects. However, the PET results were robust, consistent across participants, and localized in all the expected regions, suggesting that the statistical power allowed us to detect biologically meaningful effect sizes. For reasons outlined above (see methods), we only included female participants, which may limit the generalizability of our results to males. Finally, due to using a passive control condition (no music), the observed *BP*_ND_ changes cannot be unequivocally attributed to music-induced pleasure alone. However, the participants experienced high pleasure during the music condition, and strong *BP*_ND_ changes were observed in the reward circuits, indicating that pleasure was the main driver of the PET effects.

## Conclusions

We conclude that the endogenous opioid system plays a modulatory role in music-induced pleasure. This was evident through i) observed changes in endogenous MOR tone during pleasurable music listening in the PET experiment and ii) the ability of MOR tone to modulate pleasure-dependent hemodynamic responses in fMRI. These results underscore the involvement of MORs not only in primary reward processing but also in mediating pleasure responses to abstract, aesthetic rewards. Clinical studies should further explore whether MOR signaling mediates the analgesic, emotional, and cognitive effects of music-based interventions for pain and neuropsychiatric disorders.

## Acknowlegements

This study was supported by Academy of Finland (grant #350416)

## Supplementary Materials for

**Figure S1.**
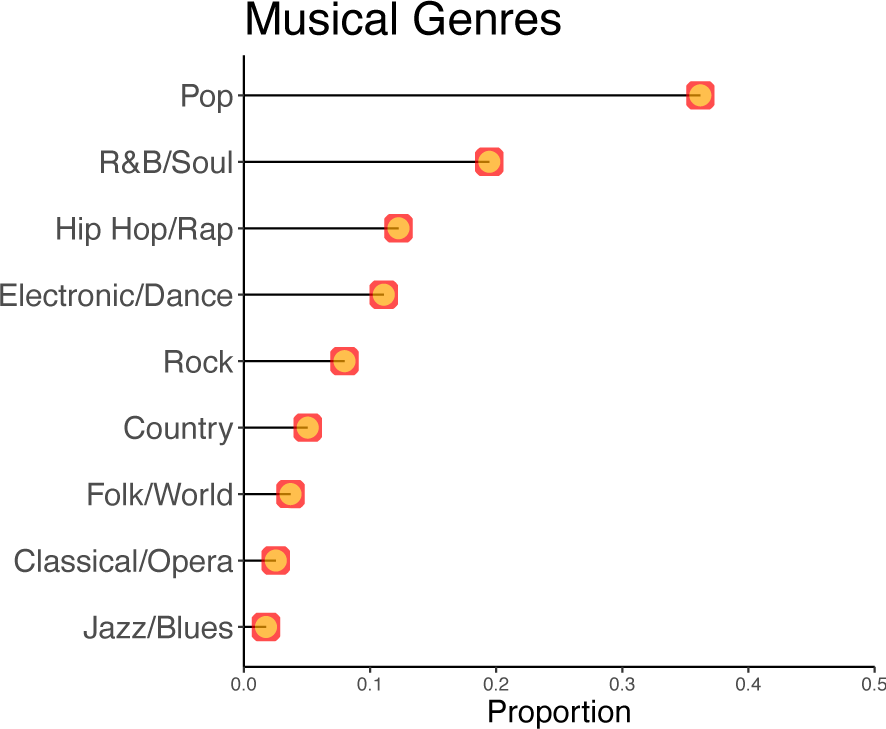
The genres for the music selected by the subjects.

**Figure S2.**
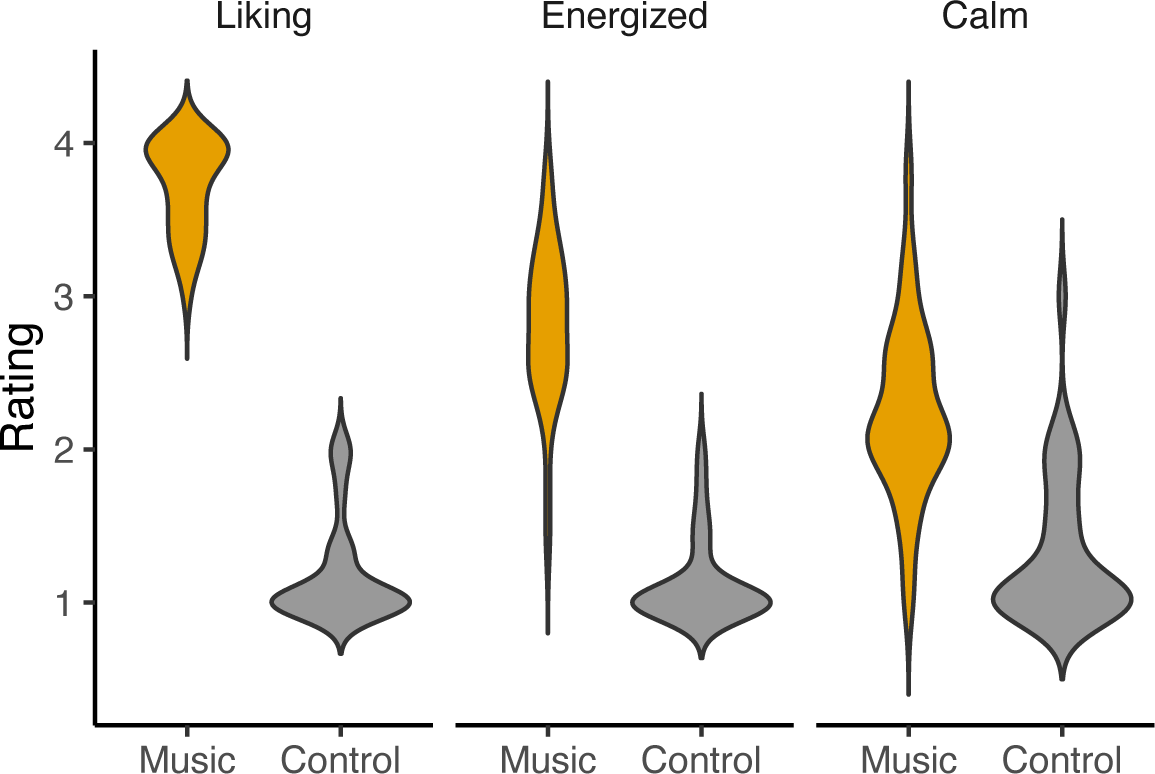
Mean ratings for liking, energization and calmness for the musical excerpts and control stimuli in the eye-tracking experiment.

